# A mammalian herbivore prefers locally adapted populations in a common garden: implications for climate change mitigation

**DOI:** 10.1101/2021.12.01.470780

**Authors:** Kayla K. Lauger, Sean M. Mahoney, Elizabeth M. Rothwell, Jaclyn P. M. Corbin, Thomas G. Whitham

## Abstract

1. Climate change is expected to alter habitat more rapidly than the pace of evolution, leading to tree populations that are maladapted to new local conditions. Assisted migration is a mitigation strategy that proposes preemptively identifying and planting genotypes that are robust to the expected climate change-induced alterations of an area. Assisted migration however, may impact the broader community, including herbivores which often coevolved with local plant genotypes and their defenses. Although this question has been examined in arthropod herbivores, few studies have assessed this question in mammalian herbivores, and fewer still have leveraged experimental design to disentangle the genetic contribution to herbivore preference.
2. We examined the hypothesis that North American porcupine (*Erethizon dorsatum*) browsing on Fremont cottonwood (*Populus fremontii*) is under genetic control in a common garden, which allowed us to uncouple genetic and environmental contributions to browse preference.
3. Generally, porcupines selected local trees and trees from climatically similar areas, where trees from local and cooler climate populations suffered over 2x more extensive herbivory than trees from warmer areas. Plant genotype was a significant factor for selection, with the most heavily browsed genotype having on average >10x more herbivory than the least heavily browsed. Because genotypes within and among populations were replicated, we calculated broad-sense heritability in which tree palatability by porcupines was H^2^_B_ = 0.28 (95% CI: 0.13-0.48) among genotypes.
4. *Synthesis and applications*. Our results indicate a genetic component to tree defenses against porcupine herbivory that can be predicted by the climate of the source population. This result has important implications for mammalian herbivores if climate change renders local tree genotypes maladaptive to new conditions. We recommend assisted migration efforts consider this implication and plant stock from both warmer and climatically similar areas to maintain genetic diversity in a changing environment, productivity and forage for mammalian herbivores.

## INTRODUCTION

Climate change is expected to alter local conditions by increasing temperatures and decreasing precipitation, resulting in natural selection pressures on tree populations to adapt to new climates (Wang et al. 2006, Hultine et al. 2020). The rate of climate change however is predicted to outpace that of evolution threatening the existence of locally adapted trees and their associated communities (Aitken et al. 2008). As an example of the severity of climate change, tree ring analyses in the American Southwest show that the ongoing droughts that began in 2000 and continue today represent a megadrought that is the second worst in 1200 years (Williams et al. 2020). To mitigate the expected decrease in forest productivity, planting trees sourced from warmer areas, known as assisted migration, is hypothesized to provide populations with adaptive genotypes capable of persisting under climate change-induced environmental disturbances (O’Neill et al. 2008, Atiken and Whitlock 2013). While trees planted via assisted migration will perform well in the future (Grady et al. 2015, Cooper et al. 2019), understanding how the broader community responds to different source populations and individual genotypes, including herbivores that rely on trees for food, remains an important question.

Previous studies that have investigated this question in arthropod herbivores have found a genetics-based preference (Bangert et al. 2006, Bangert et al. 2013) resulting in cascading community effects among trophic levels (Bailey et al. 2006, Busby et al. 2015). Studies investigating this question in mammalian herbivores have been far fewer, but have similarly shown feeding preference for genetics-based plant traits (Bailey et al. 2004, Diner et al. 2009). However the impacts of mammalian herbivore feeding preferences in the context of assisted migration (where trees from genetically distinct populations are transplanted into an herbivores range) have not been studied, despite the profound effects mammalian herbivores can have on tree health and abundances (Johnston and Naiman 1990, Durban et al. 2021), which can influence other taxa and trophic levels (Bailey and Whitham 2002, Schieltz and Rubenstein 2016, Malm et al. 2020).

North American Porcupines (*Erethizon dorsatum*) are an important group to investigate the genetic x environmental (GxE) interactions and community-wide effects of assisted migration due to their broad range across North America and their specialization for digesting tree cambium (Thornbury et al. 2019), which leads to obvious damage and can girdle trees (Olson and Lewis 1999). Porcupines may respond to assisted tree migration in several ways. First, there may be a genetic effect where porcupines are adapted to the defenses of local trees, so they may preferentially feed on trees from populations derived from sites nearest the common garden. In this scenario, palatability of trees is most related to the co-evolution of local trees and their associated herbivores and would be a measurably heritable trait in the broad sense. Under this hypothesis, porcupines, and their influence on the broader community, will persist in the short-term, but increased mortality due to preferential herbivore selection coupled with climate change-induced disturbances may reduce success of local trees. Trees from warmer climates would have an additional advantage as planting sites continue to warm, potentially causing resource shortages for porcupines. Second, trees transplanted via assisted migration may perform poorly in the short-term (Grady et al. 2015), subsequently increasing preference by porcupines. In this case, there may be a specific genetic x environment interaction in which poor adaptation to the environment of the planting site make these trees more palatable to porcupines. With this hypothesis, porcupine herbivory may jeopardize the long-term success of assisted migration efforts as they feed on trees that would perform best in the long-term. Finally, there may be no genetic effect on porcupine preferences and they feed on all populations similarly.

Here we simultaneously tested these hypotheses by leveraging an experimental Fremont cottonwood (*Populus fremontii*) common garden in Northern Arizona to assess how porcupines would respond to assisted tree migration. The use of a common garden experiment allowed us to simulate assisted tree migration by sourcing cottonwoods from genetically-distinct populations (Grady et al. 2015) along broad climatic and elevation gradients, thus subjecting local porcupines to a cafeteria-style choice experiment in a common garden setting. Results from this study are important contributions to the understanding of genetic x environmental (GxE) interactions between plants and the broader community, and how climate change, and efforts to mitigate its consequences, influence plants and their associated community members.

## METHODS

### Study site

This research was conducted in the Chevelon Creek Common Garden near Winslow, AZ (Elev. 1498m) in September, 2016. The garden is composed of Fremont cottonwood (*Populus fremontii*), velvet ash (*Fraxinus velutina*), Goodding’s willow (*Salix gooddingii*), and coyote willow (*Salix exiguia*) collected from 84 riparian populations in the Southwestern United States and Mexico (Figure 1). In each sampling location, individual trees were selected haphazardly and separated by a minimum distance of 30m to better sample the genetic diversity of a given source population. Although Fremont cottonwoods are not known to form clones (Durban et al. 2020), this spacing also avoids the possibility of collecting multiple samples of the same clone. Each population was given a population code representing the location of its source population (e.g., KC for Keams Canyon, Arizona, Table 1) and each genotype was given an identification number that follows its source population code (e.g., KC380). Of the seven populations of *P. fremontii* used in our study, six represented genotypes from the Sonoran Desert ecotype and one was from the Utah High Plateau ecotype (Ouray National Wildlife Refuge site) (Ikeda et al. 2017). Trees were tagged with their identification number using anodized aluminum tags inserted into the soil with steel wire. The trees were planted randomly in blocks generally resulting in neighboring trees from different source populations. Two blocks, hereafter “North” and “West,” were sampled in this study. All cuttings were irrigated via drip-line with 19 L of water from Chevelon Creek, three times per week during the growing season (March to November). To control for intra-garden variation in biotic and abiotic environment, we included garden block as a random effect in all models to account for this variation (see *Statistical Analyses* below).”

**Figure 1.**
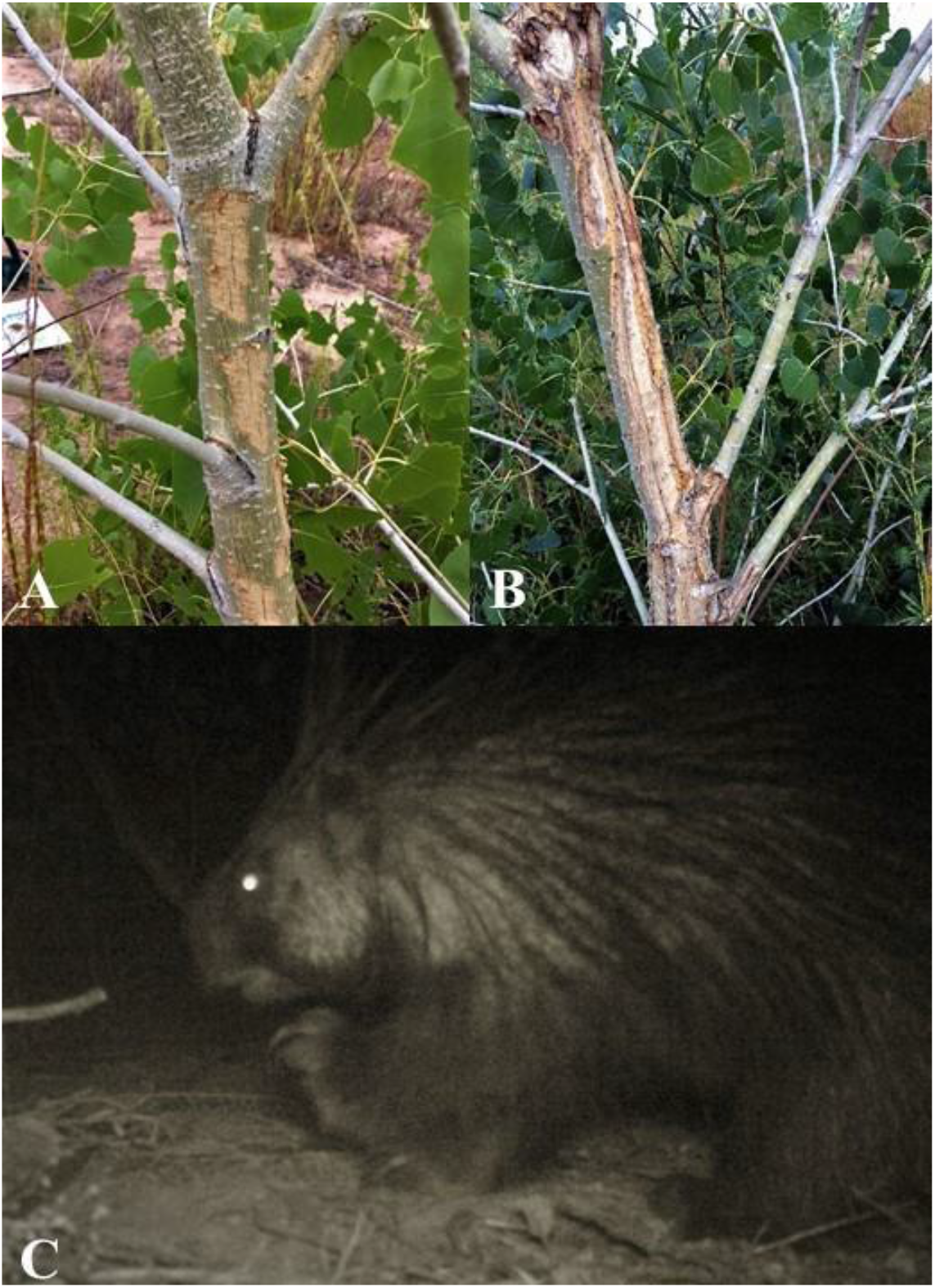
a) A “1” on the herbivory scale, typically a “nibble” or a less-than-fist-sized chunk. b) a “4” on the herbivory scale, with more than 50% of the bark on the trunk and branches being stripped. c) A North American porcupine (*Erethizon dorsatum*) captured on a trail camera set near trees with high levels of fresh herbivory.

**Table 1.** The population name, genotype name, sample sizes (n=213 total), average temperature from the warmest month over 30 years (World Clim BIO5, “MAMT”), temperature transfer (population temperature-Chevelon Common garden temperature), mean extent of porcupine herbivory (SD), mean tree stem diameter (SD) and location data for cottonwoods (*Populus fremontii*) used in study.

| Population | Population Code | State | Latitude | Longitude | Elevation (m) | MAMT | Temp. Transfer (°C) | n | Average Herbivory | Herbivory SD | Diameter (mm) | Diameter SD |
| --- | --- | --- | --- | --- | --- | --- | --- | --- | --- | --- | --- | --- |
| Ash Creek | DO | AZ | 34.40 | -112.07 | 1152 | 35.9 | -1.5 | 1 | 4.00 | - | 96.00 | - |
| Beaver Creek | BCSE | AZ | 34.67 | -111.71 | 1143 | 36 | -1.9 | 1 | 1.00 | - | 89.80 | - |
| Calf Creek | CALF | UT | 37.79 | -111.41 | 1624 | 32.9 | 1.5 | 6 | 1.17 | 0.75 | 63.55 | 19.10 |
| Cibola National Wildlife Refuge | CR | AZ | 33.36 | -114.70 | 70 | 42.1 | -7.7 | 19 | 1.58 | 1.69 | 65.92 | 27.15 |
| Citadel Wash | CW | AZ | 35.61 | -111.32 | 1299 | 36.5 | -2.1 | 1 | 2.00 | - | 33.40 | - |
| Fremont River | FR | UT | 38.28 | -111.09 | 1473 | 34.9 | -0.5 | 3 | 0.25 | 0.35 | 75.65 | 19.73 |
| Gila River | GR | AZ | 32.97 | -109.31 | 1048 | 37.7 | -3.3 | 26 | 0.05 | 0.12 | 58.13 | 14.61 |
| Globe Seven Mile | GSM | AZ | 33.58 | -110.65 | 1218 | 15.7 | -0.2 | 1 | 0.00 | - | 94.40 | - |
| Jack Rabbitt | JR | AZ | 34.96 | -110.44 | 1507 | 34.4 | 0 | 26 | 1.42 | 0.95 | 68.83 | 15.40 |
| Keam's Canyon | KC | AZ | 35.81 | -110.17 | 1920 | 31.1 | 3.3 | 50 | 1.33 | 1.00 | 64.56 | 13.07 |
| Kirkland Ash Creek | KIRK | AZ | 34.45 | -112.80 | 1555 | 35.1 | -0.7 | 15 | 0.38 | 0.88 | 67.31 | 14.33 |
| Lake Mary | LM | AZ | 35.07 | -111.53 | 2109 | 27.7 | 6.7 | 12 | 2.06 | 1.47 | 77.28 | 10.85 |
| Mount Meadow Massacre Memorial | MMM | UT | 37.48 | -113.64 | 1726 | 30.7 | 3.7 | 2 | 0.50 | 0.71 | 72.15 | 20.58 |
| Muley Twist Capital Reef | MTCR | UT | 37.85 | -111.03 | 1730 | 32.3 | 2.1 | 2 | 1.50 | 0.71 | 64.50 | 23.19 |
| North Cottonwood Creek | NCC | UT | 38.08 | -109.57 | 1624 | 33.7 | 0.7 | 23 | 1.15 | 0.84 | 67.13 | 13.40 |
| Ouray National Wildlife Refuge | OR | UT | 40.15 | -109.62 | 1423 | 34.2 | 0.2 | 20 | 1.50 | 0.63 | 67.05 | 11.20 |
| Palo Verde Ecological Reserve | PV | AZ | 33.70 | -114.50 | 87 | 42.2 | -7.8 | 1 | 0.00 | - | 86.30 | - |
| Skull Valley | SV | AZ | 34.53 | -112.71 | 1346 | 33.3 | 1.1 | 3 | 2.67 | 2.31 | 57.60 | 7.84 |
| Trachete Creek | TRCH | UT | 37.96 | -110.57 | 1439 | 35.3 | -0.9 | 1 | 2.00 | - | 55.80 | - |
| Chevelon Experimental Common Garden | CHEV | AZ | 34.99 | -110.64 | 1507 | 34.4 | 0 | - | - | - | - | - |

### Herbivore identification

We set a motion sensor Ambush^®^ camera in the garden on two separate occasions, once in the North block and once in the West block. In both cases, the camera faced trees that were freshly and extensively browsed. The cameras were activated for a total of 276 hours.

### Sampling and data collection

To obtain an accurate representation of the garden, we visually inspected each tree (n=213, Table 1) in the North and West blocks of the Chevelon Common Garden for evidence of mammalian herbivory in the form of chew marks on the bark. Specifically we quantified tree damage to the base, trunk, nodes, and branches, visually starting at the base and travelling up to the canopy of the tree. The extent of herbivory was classified on a 0-4 scale, with 0 representing no damage, 1 representing little damage (such as a “nibble”; Figure 2a) and 4 being extensive bark-stripping of >50% of the trunk and branches (Figure 2b).

**Figure 2.**
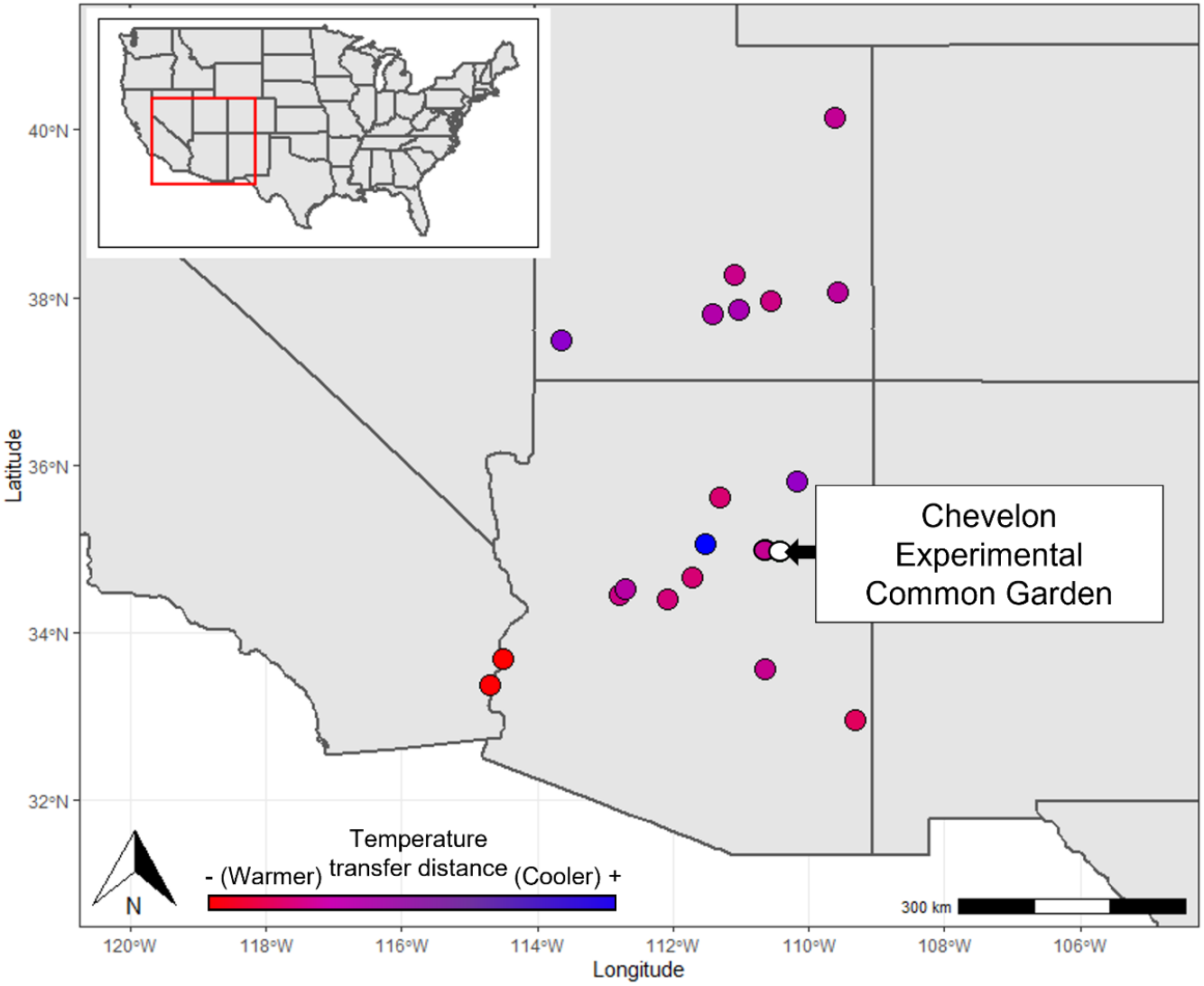
A map of the geographical locations of source populations planted in the Chevelon Common Garden. Red dots indicate warmer, while blue dots indicate cooler source populations relative to the Chevelon Common Garden (white dot).

We also assessed the performance of each tree by measuring tree diameter and classified each tree as either shrub-like or tree-like. A previous study at Chevelon Garden has shown that poorly performing trees were shorter, had smaller trunks, and had more stems (i.e., “shrub-like”) than better performing trees (Mahoney et al. 2018). The architecture of each tree was classified as either “shrub” or “tree” with shrubs having more than two trunks. The diameter at the stem base (DSB) was measured for all trees, the largest trunk was measured for trees with a “shrub” architecture.

### Temperature transfer distance

To assess the relationship between temperature transfer distance and porcupine herbivory, we estimated the mean annual maximum temperature (MAMT) of each population and the Chevelon common garden using World Clim maximum temperature of the warmest month (BIO5) recorded during each year averaged over 30 years at 30 seconds (approximately 1 km^2^) resolution (Table 1). We then calculated the temperature transfer distance for each population as the difference between the temperature at Chevelon and the temperature of the population (Chevelon_MAT_ - Population_MAT_ = temperature transfer distance; Grady et al. 2011, Mahoney et al. 2018, Cooper et al. 2019). Temperature transfer distance quantifies changes in local climatic conditions, including elevation and latitude, and is represented as the difference in maximum temperature (either positive or negative) between the common garden and the source population home site. A negative temperature transfer represents trees from a warmer source population, while a positive temperature transfer represents trees from a cooler climate (Mahoney et al. 2018).

### Statistical analysis

We assessed our hypothesis that porcupine herbivory was related to temperature transfer distance in several ways. First, we used a generalized linear mixed effects model with binomial distribution to test if tree herbivory was related to temperature transfer distance. In this analysis, herbivory was the response variable and was treated as a binomial variable (yes [n=111] or no [n=102], total n=213). Population temperature transfer distance, stem diameter, and architecture type (shrub-like or tree-like) were main effects, and garden quadrant (North or West block) was included as a random effect. In this model, a tree was categorized as being affected by herbivory if any amount of porcupine herbivory was evident.

Next, we tested whether the extent of tree herbivory was related to temperature transfer using a generalized linear mixed effects model with a Poisson distribution. We used a Poisson distribution in this analysis because our estimates of porcupine herbivory were discrete counts. In this model, extent of herbivory was the response variable, population temperature transfer, stem diameter, and tree architecture were the main effects, and the garden quadrant was included as a random effect.

We then assessed if porcupine herbivory varied among populations and specific tree genotypes using separate linear mixed effects models. In the model testing the population effect, the extent of herbivory was the response variable, population, stem diameter, and architecture type were the main effects and garden quadrant was the random effect. In the model testing the plant genotype effect, the extent of herbivory was the response variable, plant genotype, stem diameter, and architecture type were main effects and garden quadrant was the random effect. Only populations with n≥3 trees (any genotype) and tree genotypes with n≥3 replicates were included (Table 1). We assessed model assumptions by inspecting QQ and residual plots and all analyses were conducted in R using lme4 package.

Finally, we calculated the broad-sense heritability (H^2^_B_) of tree palatability as the variation in the phenotype of the browsed plants, (r2s), divided by the total variance in the phenotype for all genotypes (r2 total), or H^2^_B_ = r2s/r2 total (Shuster et al. 2006). Our heritability calculations used an average of 5 replicates per genotype and genotypes with less than 3 sampled individuals were omitted from the analysis. Heritability was estimated for a subset of genotypes within the garden and is presented as H^2^_B_ ± 95% confidence intervals. The intervals were found by multiplying the standard error by the t-scores matching sample sizes for each population. As such, the effects of genotype on porcupine preference would not be statistically significant if the 95% CI overlaps with 0. All analyses were completed in R using the Heritability package developed by Kruijer et al. (2015).

## RESULTS

### Herbivore identification

The North American porcupine (*Erethizon dorsatum*) is the most likely agent for the type and extent of herbivory that we have observed. Xylophagous rodents captured on the trail camera (Figure 2c) and observed during data collection included porcupines, and ground squirrels. While bark stripping up to the canopy of a tree has been documented in both porcupines and ground squirrels (Scheffer 1952), diagonal climbing marks and clumps of large chew marks indicate porcupine herbivory (Diner et al. 2009).

### Herbivore preferences – temperature transfer distance, population, genotype

From our logistic regression, in which trees were classified as either damaged or not, trees from populations of similar climate to Chevelon and trees with positive MAMT transfer distances (i.e., trees from cooler climates), were more likely to exhibit porcupine herbivory (Figure 3, **β**=0.26, z=4.3, p<0.0001, garden quadrant ***σ***^2^=3.6). Diameter did not explain the likelihood of herbivory (**β**=0.002, p=0.33).

**Figure 3.**
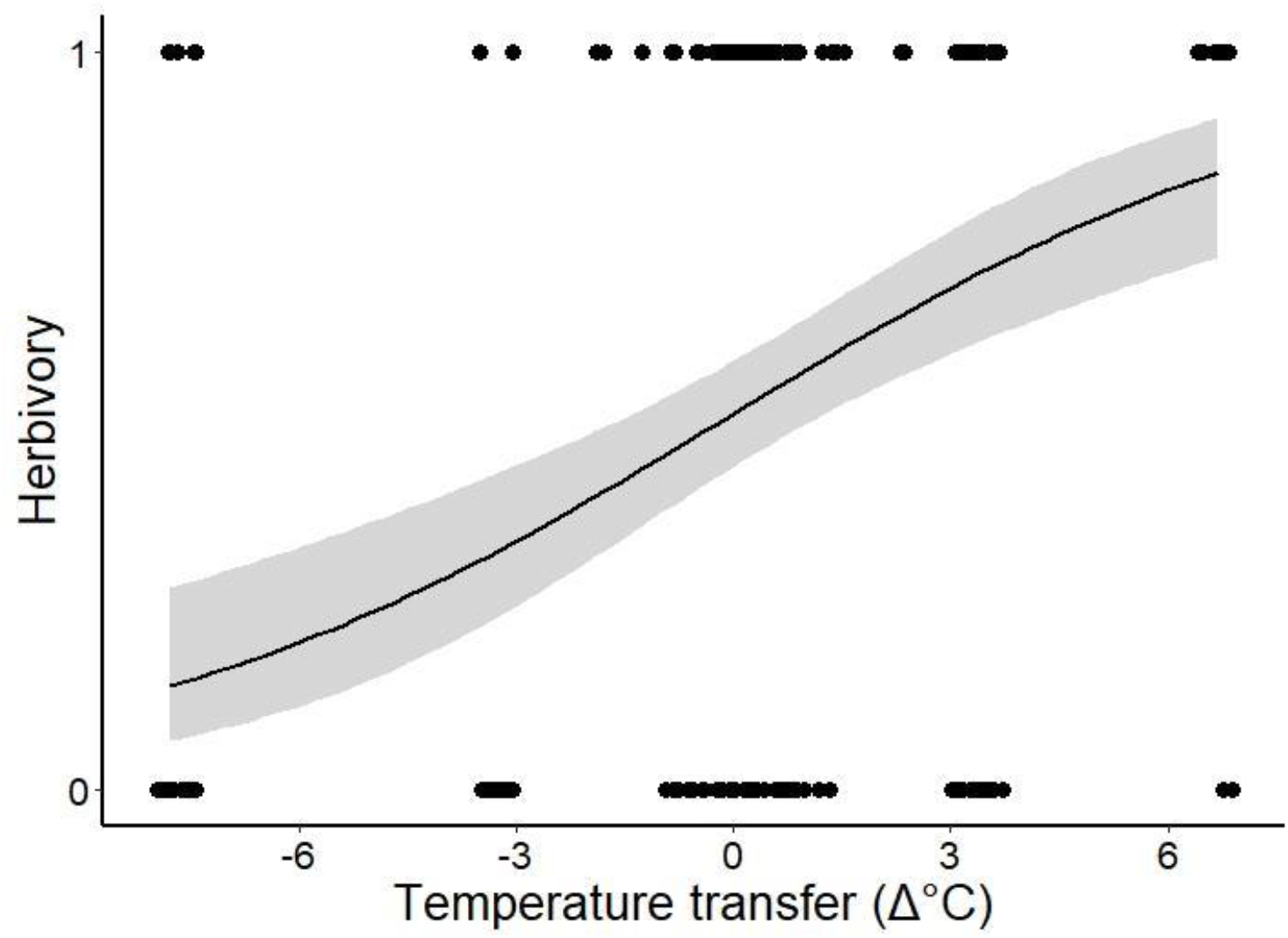
Trees from populations that are climatically similar to or cooler than the Chevelon Common Garden were more likely to exhibit porcupine herbivory (logistic regression: **β**=0.26, p<0.0001). In this analysis, tree were classified as either damaged or not, and the curve represents the probability ±SE of a tree exhibiting herbivore damage based on its temperature transfer distance.

The generalized linear model of our herbivory classifications (0-4) showed that the extent of herbivory increased with MAMT, where trees from warmer climates experienced less extensive herbivore damage than trees from climates similar to Chevelon (Figure 4, **β** =0.1, p<0.0001, garden quadrant ***σ*** ^2^=0.09). Trees that were shrub-like in architecture experienced less extensive herbivory damage (Figure 5, **β** =0.4, p=0.01).

**Figure 4.**
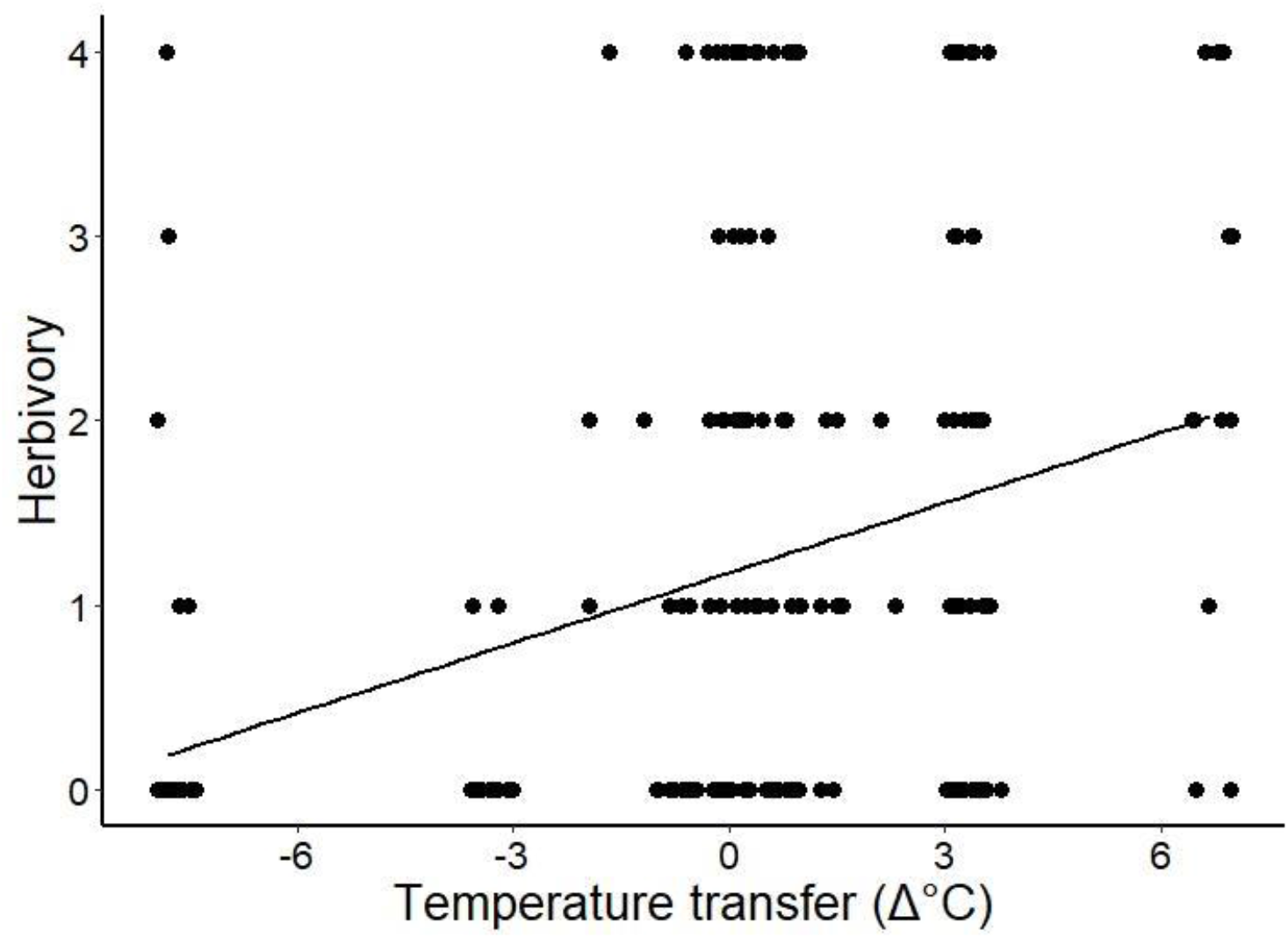
Trees from populations that are climatically similar to or cooler than the Chevelon common garden experienced more extensive porcupine herbivory (generalized linear model: **β**=0.1, p<0.0001). Herbivore score is represented by discrete classifications based on visual inspection of each tree. An herbivore score of 0 represents no damage present, while 4 indicated extensive herbivore damage (see Figure 1 for example photographs).

**Figure 5.**
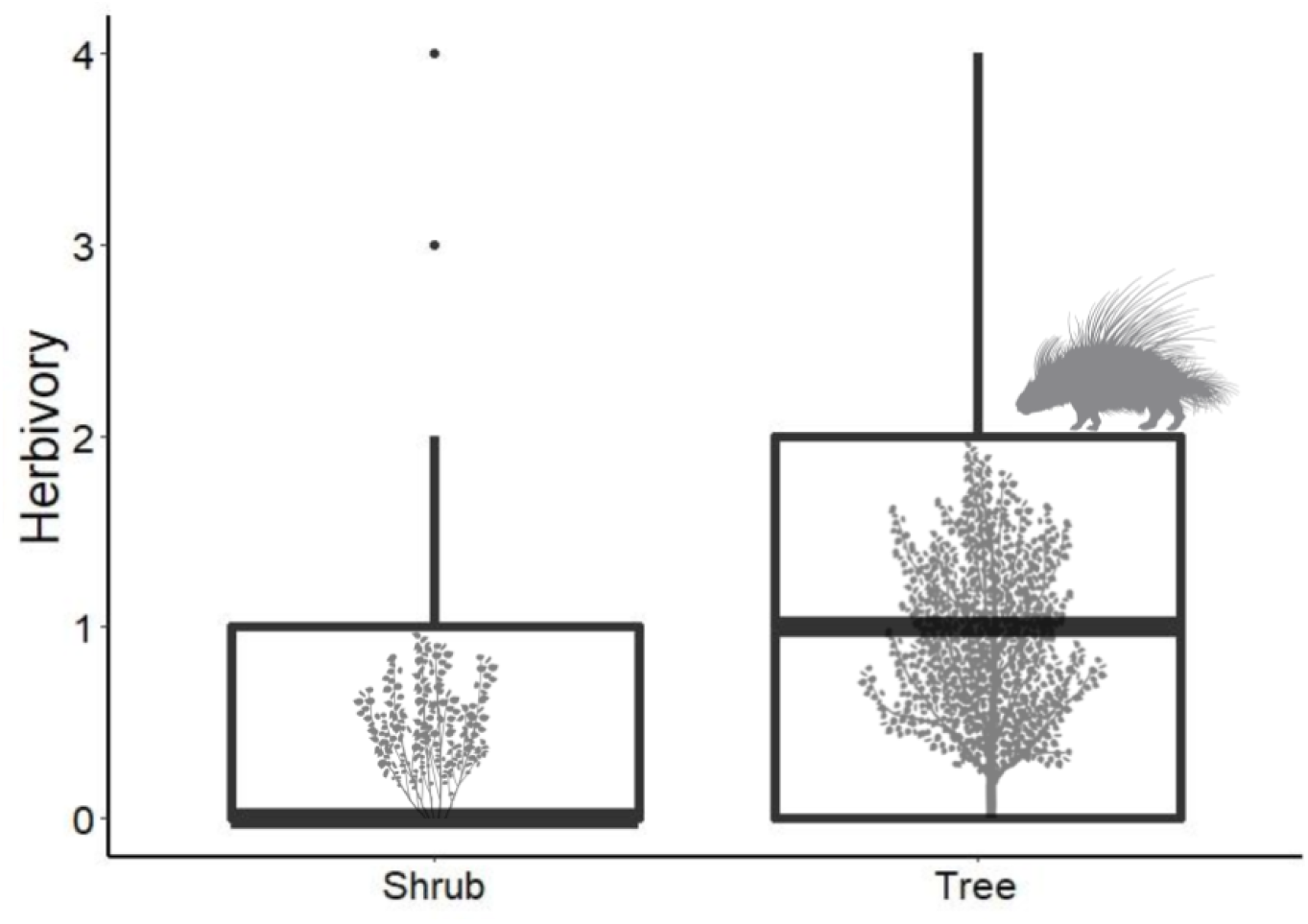
Trees that were shrub-like in architecture experienced less herbivory damage (generalized linear model: **β**=0.4, p=0.01). Herbivore score is represented by discrete classifications based on visual inspection of each tree. An herbivore score of 0 represents no damage present, while 4 indicated extensive herbivore damage (see Figure 1 for example photographs). Boxplots indicate 1st and 3rd quartiles (boxes), median (thick horizontal line), range (whiskers) and outliers (dots).

From our linear mixed effects models, we found that populations were differentially affected by porcupine herbivory (F_10,186.21_=3.4, p=0.0004, garden quadrant ***σ*** ^2^=0.66). This effect was largely driven by differences between population GR and sites from intermediate and cooler transfer distances, where trees from GR experienced less herbivory (Figure 6, posthoc comparisons between GR and JR (t=-4.0, p=0.004), KC (t=-3.9, p<0.007), LM (t=-3.3, p<0.05), OR (t=-3.6, p<0.02), and SV (t=-3.5, p<0.03). All other comparisons were not different.

**Figure 6.**
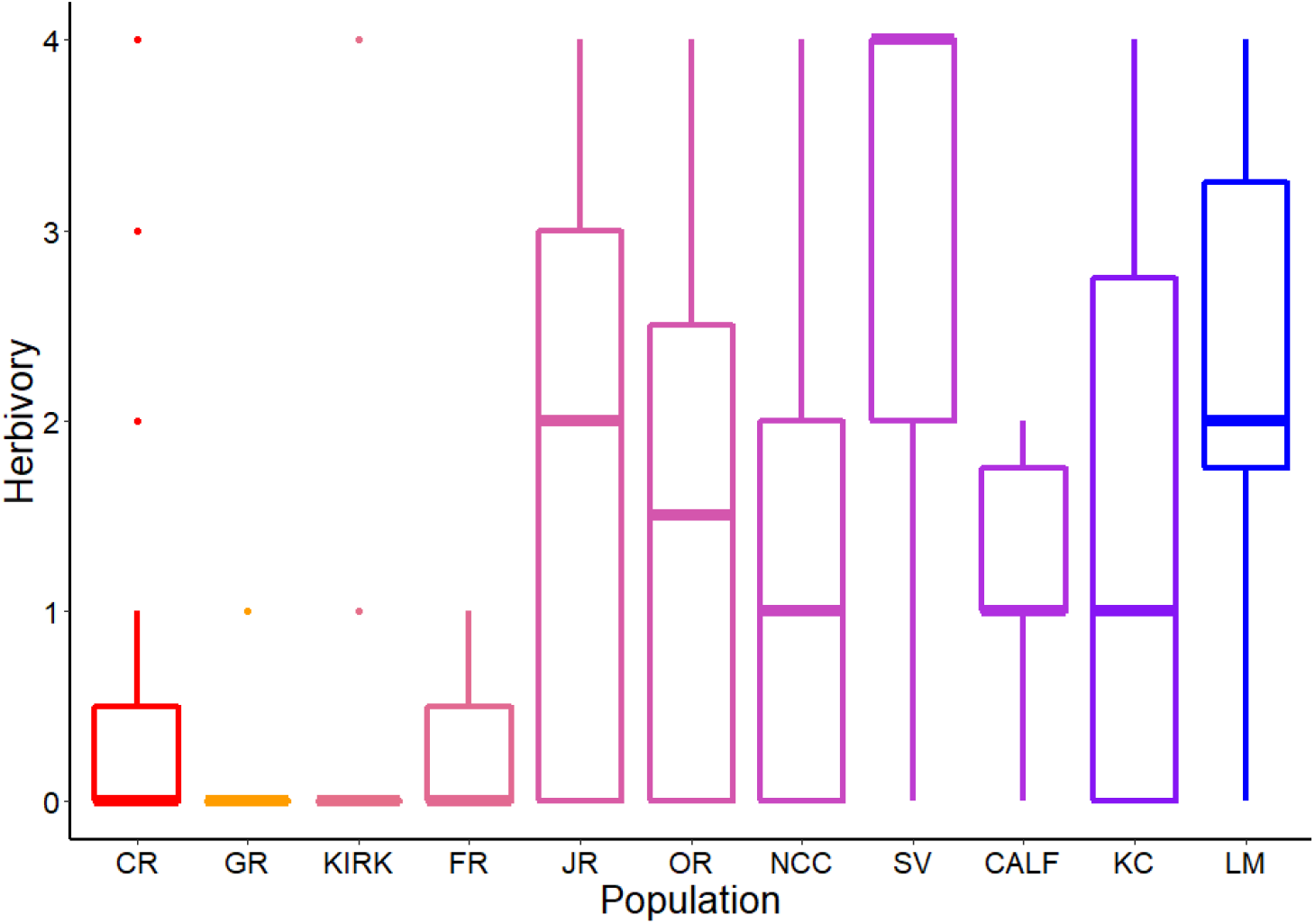
Tree populations were differentially affected by porcupine herbivory (mixed effects model: F_10,186.21_=3.4, p=0.0004). Herbivore score is represented by discrete classifications based on visual inspection of each tree. An herbivore score of 0 represents no damage present, while 4 indicated extensive herbivore damage (see Figure 1 for example photographs). Red boxes indicate warmer, while blue dots indicate cooler source populations relative to the Chevelon Common Garden. Boxplots indicate 1st and 3rd quartiles (boxes), median (thick horizontal line), range (whiskers) and outliers (dots).

Generally, the extent of porcupine herbivory differed among plant genotypes (F_26,49.2_=2.82, p=0.0009, garden quadrant ***σ***^2^=0.48). Specifically, one genotype from the KC population suffered greater herbivory than genotypes from CR and GR. From posthoc comparisons, KC380 exhibited significantly more herbivory than CR86 (t=-4.2, p<0.02), GR271 (t=-4.6, p<0.003), GR276 (t=-4.3, p<0.01), and GR278 (t=-4.5, p=0.004). OR7 exhibited marginally more herbivory than GR271 (t=-3.7, p=0.08). Comparisons among all other genotypes were not significant.

### Heritability of palatability

Estimations of broad-sense heritability (H^2^_B_) revealed that the palatability of genotypes is a heritable trait. At the Chevelon garden, genetic variation explained H^2^_B_ = 0.28 (95% CI: 0.13-0.48) of the variation in porcupine preference at the genotype level. Among genotypes within a single common garden block, we also found that the estimated genetic variance is 0.07, with a residual variance of 0.18. Heritability assessed at the population level revealed a heritability estimate of H^2^_B_ = 0.31 (95% CI: 0.13-0.67). Among populations the genetic variance is 0.08 with a residual variance of 0.18. We detected both a population and genotype effect on heritability, with population having a greater estimate of heritability which is likely due to the increased number of samples at the population level compared to the genotype level. These findings suggest that palatability for porcupine preference is a heritable trait in the broad-sense (i.e., clonal) and that local genotypes are innately more preferred than others in a large-scale cafeteria context.

## DISCUSSION

Our results indicate MAMT transfer distance of cottonwood source populations has a pronounced effect on susceptibility to mammalian herbivory, supporting a genetic component to herbivore defense. At the Chevelon Common Garden, a relatively high elevation planting site, trees from similar, or cooler MAMT populations exhibited approximately 2-3x more extensive damage from porcupines than trees from warmer MAMT sites. Our findings support the hypothesis that tree herbivore defense is genetics-based and imply that porcupines prefer to browse on trees from source populations with similar climate, or trees that are adapted to the local climate of the garden (Diner et al. 2009, Tomas et al. 2011, Holeski et al. 2012, Stone et al. 2018). Interestingly, with porcupines being more abundant in the northern portion of the state than in the south, the distribution of porcupines in Arizona generally correlates with the locations of populations that were heavily browsed in this study (Brown and Babb 2009). This corroborates our results, suggesting that porcupines are: 1. adapted to the defenses of the trees they encounter most often, 2. that trees from cooler climates do not place as much energy into creating phytochemical defenses, 3. trees from warmer climates are poorer performers when planted in cooler climates and lack sufficient nutrition such that they are avoided or not recognized as a resource, or 4. some combination of the above.

There were significant differences in the extent of herbivory between plant genotypes from different source populations. Genotype KC380 showed significantly more evidence of herbivory than CR86 and three genotypes from the GR population. While trees from the KC population experienced relatively high levels of herbivory and those from the GR population experienced relatively low levels overall, significant differences among certain genotypes from these populations suggest that there may be genetically driven differences in phytochemical defenses that may be independent of climatic adaptations. Although we do not have estimates of phytochemicals from trees at Chevelon, numerous studies show plant phytochemical defenses are genetics-based, and that genotype is a significant factor affecting herbivore selection in *Populus* spp. (e.g., Martinsen et al. 1998, Wimp et al. 2007, Diner et al. 2009, Holeski et al 2012). Our study is the first we are aware of that investigates GxE interaction effects on mammalian herbivory in the context of assisted migration, more investigation is needed to assess the role of how plant chemical defenses mediate mammalian herbivory following assisted migration.

Our data also shows that cottonwood palatability for porcupines is a heritable trait. Although we didn’t investigate the underlying mechanism(s) in our study, Diner et al. (2009) found that two phenolic glycosides (tremulacin and salicortin), which are under strong genetic control were the chemical variables most associated with feeding choices by porcupines in wild stands of aspen (*P. tremuloides*). Bark traits may also be important to consider with cottonwoods; for example, in common garden studies, Lamit et al. (2015b) found that bark roughness was a highly heritable trait (H^2^_B_ = 0.48), which in turn played a major role in defining the lichen community on the tree trunk. With other mammalians such as the North American beaver (*Castor canadensis*), Bailey et al. (2004) used cafeteria studies in which branches from different tree genotypes were placed around beaver ponds. They found that beavers selectively avoided cottonwood tree genotypes high in condensed tannins (Bailey et al. 2004). Similarly, Bailey et al. (2007) found that elk (*Cervus elaphus*) herbivory on aspen (*Populus tremuloides*) was negatively associated with tremulacin concentration. Lastly, Bailey et al. (2006) found that trophic interactions were associated with cottonwood plant chemistry (condensed tannin and salacortin), susceptibility to gall aphids (*Pemphigus betae*), and the foraging of insectivorous birds in which the tri-trophic interaction was quantified as a heritable plant trait (H^2^_B_ = 0.62 ± 0.24). Although relatively few studies have examined the genetic basis of mammalian generalist foraging behavior, it appears that plant chemistry has played an important role in medium to large mammals and that their selectivity for some genotypes and populations over others could feedback to affect plant evolution.

Preferential mammalian herbivory can foil plant success and act as a genetic-based selection pressure for climatic adaptation (Cooper et al. 2019) or other genetics-based traits by browsing trees to the point of girdling and death. This not only has implications for survival of tree populations through climate change, but can lead to cascading effects in community structure and the evolution of whole communities (Whitham et al. 2012, 2020). Previous studies have shown community-level effects of selection by arthropod herbivores acting on individual genotypes and populations (Keith et al. 2010) leading to much larger effects on communities of diverse organisms (Busby et al. 2015, Lamit et al. 2015a). Mammalian herbivores can also have large impacts on success through selection that changes the underlying genetic structure of natural populations and communities (e.g., rapid evolution by goldenrod and their associated arthropod communities in response to elk browsing; Smith et al. 2015).

Our results lend strong support to the hypothesis that selection by mammalian herbivores is genetically based, and that herbivore populations are adapted to their local plant populations (Bailey et al. 2004, Durban et al. 2021). Other studies from Chevelon showed local genotypes were more resilient to fungal infections and invasive plant competitors than genotypes sourced from areas >3°C warmer (Grady et al. 2015, Mahoney et al. 2018). This presents an added challenge to managers because, in the face of climate change where increased temperatures may exceed 3°C, non-local genotypes from areas >3°C warmer will need to be planted for long-term gains, however in the short-term these maladaptive genotypes will suffer from relatively low productivity. Consequently, local genotypes will also need to be planted to maintain habitat in the short-term and, as our results clearly show, support local mammalian herbivore communities. Therefore, as with previous recommendations, to meet the short- and long-term challenges of climate change-induced environmental disturbances (Seager et al. 2007, Garfin et al. 2013, Wuebbles et al. 2017), a mix of local and non-local stock from areas up to 3°C warmer should be selected for assisted migration efforts (Grady et al. 2015, Mahoney et al. 2018, Cooper et al. 2019).

## ACKNOWLEDGEMENTS

We thank Arizona Game and Fish Department and Arizona State Forestry Division for financial support in establishing and maintaining the common garden. We thank Karla Kennedy and Kevin Grady who played important roles in the development and establishment of the common garden and T. C. Theimer who provided game cameras.

## AUTHORS’ CONTRIBUTIONS

K.K.L and E.M.R conceived of project, designed methodology, collected all field data, and wrote initial manuscript. S.M.M analyzed data. J.P.M.C analyzed data and contributed to acquisition of metadata. T.G.W. contributed to study design, conceptual framework and obtained funding for the establishment of the common garden. All authors contributed to writing the revised manuscript. Tree schematics (Figure 5) drawn by J.B. Mike.

